# Garlic-derived organosulfur compounds regulate metabolic and immune pathways in macrophages and attenuate intestinal inflammation in mice

**DOI:** 10.1101/2021.11.02.466911

**Authors:** Ling Zhu, Laura J. Myhill, Audrey I.S. Andersen-Civil, Stig M. Thamsborg, Alexandra Blanchard, Andrew R. Williams

## Abstract

**Scope:** Garlic is a source of bioactive phytonutrients that may have anti-inflammatory or immunomodulatory properties. The mechanism(s) underlying the bioactivity of these compounds and their ability to regulate responses to enteric infections remains unclear.

**Methods and Results:** We investigated if a garlic-derived preparation (PTSO-PTS) containing two organosulfur metabolites, propyl-propane thiosulfonate (PTSO) and propyl-propane thiosulfinate (PTS), regulated inflammatory responses in murine macrophages and intestinal epithelial cells (IEC) *in vitro,* as well as in a model of enteric parasite-induced inflammation. PTSO-PTS decreased lipopolysaccharide-induced secretion of TNFα, IL-6 and IL-27 in macrophages. RNA-sequencing demonstrated that PTSO-PTS strongly suppressed pathways related to immune and inflammatory signaling. PTSO-PTS induced the expression of a number of genes involved in antioxidant responses in IEC during exposure to antigens from the parasite *Trichuris muris. In vivo,* PTSO-PTS did not affect *T. muris* establishment or intestinal T-cell responses but significantly altered caecal transcriptomic responses. Notably, a reduction in *T. muris*-induced expression of *Tnf, Saa2* and *Nos2* was observed.

**Conclusion:** Garlic-derived organosulfur compounds exert anti-inflammatory effects in macrophages and IEC, and regulate gene expression during intestinal infection. These compounds and related organic molecules may thus hold potential as functional food components to improve gut health in humans and animals.

## 1. Introduction

The role of dietary components in gut health and immune function has attracted considerable attention in recent years. Plant-derived compounds, such as polyphenols, inulin and allicin, have been extensively studied for their potential roles in preventing inflammation and infectious diseases.^[1],[2]^ Moreover, the use of these phytonutrients in animal production has demonstrated several protective effects against numerous disorders by regulating the immune system or improving other properties.^[3]^ Among these, garlic *(Allium sativum),* which has been used as a form of traditional medicine and food additive, has attracted increasing interest due to its antioxidant, antimicrobial and antifungal activities.^[4]^

Garlic extracts or garlic-derived compounds can produce immunomodulatory properties by regulating the secretion of inflammatory cytokines such as IL-6, TNFα, and IL-1β, indicating its capacity to affect immune homeostasis.^[5]^ Consistent with this, garlic consumption can activate γδ-T and NK cells, resulting in fewer cold symptoms and reduced flu severity, suggesting stronger immune function and lower inflammation *in vivo*.^[6]^ The balance between T helper 1 and 2 (Th1 and Th2) responses, which are important for the adaptive immune system in terms of cytokine release and disease resistance, can be modulated in favour of a Th2 polarized response in PHA-stimulated spleen lymphocytes treated with garlic supplement.^[7]^ The distinctive properties of garlic are related to active compounds, mainly including sulfur compounds, like allicin also called diallyl thiosulfinate, and non-sulfur compounds such as phenols, flavonoids and saponins.^[8,9]^ Most studies have focused on sulfur-containing compounds, to which the flavourful and medicinal properties of garlic are attributed. For example, propyl-propane thiosulfonate (PTSO) and propylpropane thiosulfinate (PTS), garlic derived organosulfur compounds, are used as biopackaging materials for food due to their antimicrobial activity leading to enhanced food safety and longer shelf life.^[10]^ Extracts containing PTSO and PTS has also been reported to affect the intestinal mucosal thickness, and ileal villus morphology, which may be beneficial during *Salmonella* spp and *Escherichia coli* infection in broiler chickens.^[11]^ Additionally, evidence from mouse studies with PTSO-PTS mixture treatment (80% PTSO, 20%PTS) indicates PTSO-PTS may directly interact with gastrointestinal cells to regulate mucosal immune response by upregulating *Reg3g, IFNg* and *Il33* expression. ^[12]^ Furthermore, intestinal inflammatory responses induced by 2,4-dinitrobenzene sulfonic acid (DNBS) and dextran sodium sulfate (DSS) can be ameliorated with PTSO, which is likely related to the regulation of intestinal epithelial barrier as well as the modulation of intestinal microbiota.^[13]^ The protective activities of PTSO-PTS on inflammation suggests that PTSO-PTS may be used as a natural therapy against infective diseases but further investigations are still needed to understand the underlying mechanisms.

Intestinal pathogens such as parasites may cause dysbiosis, chronic inflammation, and altered gut function and nutrient metabolism.^[14]^ The murine whipworm *Trichuris muris* is a natural gastrointestinal parasite of mice, and is a valuable experimental model for the closely related human parasite *T. trichiura*.^[15]^ Low doses of *T. muris* eggs in susceptible hosts induce a chronic infection characterized by a Th1 polarized cellular response, with increased expression of cytokines such as IFNγ and IL-27. Thus, this model is a useful experimental system to study how dietary components may regulate intestinal immune function and inflammatory diseases in humans and animals.^[16]^

We hypothesized that PTSO-PTS can exert protective effects on intestinal inflammation during chronic enteric parasite infection, with the potential effects related to immune cell or intestinal epithelial cell modulation. We show that PTSO-PTS decreases cytokine release from murine macrophages, and regulates gene expression in macrophages and intestinal cells with or without either lipopolysaccharide (LPS)-stimulation or activation with *T. muris* antigens. Furthermore, the immunoregulatory properties of PTSO-PTS *in vivo* were evaluated using transcriptomic analysis of murine caecal tissue derived from *T. muris*-infected mice. Our results shed light on how PTSO-PTS regulates immune function and inflammatory responses, and encourage assessment of how compounds derived from garlic or related plants may be used to modulate enteric disease.

## 2 Experimental Section

### 2.1 Garlic extract

The garlic extract was provided by Pancosma SA (Rolle, Switzerland). It is standardized to 40% PTSO-PTS (of which 80% consists of PTSO, and 20% PTS). The remainder of the extract consists of the solvent polysorbate-80. The stated experimental concentrations refer to final PTSO-PTS concentration used in assays or animal experiments. In all experiments, control cells or mice were treated with an equivalent amount of polysorbate-80 as the PTSO-PTS-treated groups.

### 2.2 Parasites

*T. muris* parasites (strain E) were maintained and *T. muris* excretory/secretory (E/S) products prepared as described previously.^[17]^ Eggs were purified from the faeces of mice and embryonated for at least two months before being used for infection. Experimental mice were infected with 20 eggs suspended in water by oral gavage.

### 2.3 Cell culture

Murine rectal epithelial CMT93 (ATCC-CCL-223) cells or RAW264.7 macrophages (ATCC-TIB-71) were cultured in DMEM (Sigma Aldrich, USA) medium with 10% fetal calf serum (Sigma Aldrich), 100 U/mL penicillin and 100 μg/mL streptomycin (Sigma Aldrich). The cell culture was maintained at 37 °C with 5% CO2. Cell viability was tested with prestoBlue cell viability reagent (Thermo Fisher Scientific) according to the manufacturer’s protocols. Briefly, a concentration of 1×10^5^ cells per mL were seeded into 96 well plate (Sigma Aldrich). After 24h, the supernatant was removed and new medium was added into cells with 90 uL DMEM and 10 uL PrestoBlue reagent per well. The cells were incubated for further 2h then the absorbance at 490 nm with a reference wavelength at 600□nm for PrestoBlue was measured.

### 2.4 Cell stimulation

RAW264.7 cells were seeded into a 24-well plate and 2h allowed for attachment. Following this, cells were stimulated with 20 μM PTSO-PTS and 500 ng/mL LPS for either 6h or 24h. RNA was extracted from RAW264.7 cells cultured for 6h using RNeasy kits according to manufacturer’s protocols (Qiagen, Denmark). After 24h culture, supernatant from cells was collected for further analysis. For CMT93 cells, 20 μM PTSO-PTS and *T. muris* E/S (50 μg/mL) were added for 6h when cell confluence reached to 80%. RNA extraction was conducted as for RAW264.7 cells.

### 2.5 ELISA

Levels of secreted IL-6, TNFα and IL-27 were measured in RAW264.7 supernatants using Mouse paired ELISA kits (Duosets, R and D Systems, UK) according to the manufacturer’s instructions.

### 2.6 Animals and Experimental design

Female C57BL/6JOIaHsd mice (6-8 weeks of age, Envigo) were randomly divided into 4 group(n=5) and housed in ventilated cages with free access to standard murine diet (D30, SAFE) and water. Two untreated groups were included with vehicle water, the rest of groups received PTSO-PTS water. PTSO-PTS treatment commenced 7 days before infection and continued throughout the course of infection (35 days). Water was refreshed every 2 days. Mouse body weights were recorded weekly throughout all the experimental period. After one week of PTSO-PTS intake, mice were infected with *T. muris,* and were sacrificed at 35 days post infection. Caecal tissues were collected in RNAlater (Thermo Fisher) and stored at −20 °C for further RNA extraction. RNA was extracted using miRNeasy mini kits after homogenization using a gentle MACS Dissociator (Miltenyi Biotec) in QIAzol lysis buffer. This study was approved by the Danish Animal Experimentation Inspectorate (Licence number 2015-15-0201-00760) and conducted at the Experimental Animal Unit, University of Copenhagen.

#### 2.6.1 Dosage Information

Water was prepared by solubilizing PTSO-PTS to deliver a concentration 0.1 mg PTSO-PTS /kg bodyweight (BW)/day or 1 mg/kg BW/day, assuming a water intake of 4 ml/day per mouse, and control water was normal drinking water with the same volume of polysorbate 80 alone.

### 2.7 Mesenteric lymph nodes (MLN) cells isolation

MLN tissues were collected from mice at 35 days post infection, then placed into 70 μM cell strainer for tissue disruption using a sterile syringe plunger to obtain a single cell suspension. RPMI media (Life Technologies) with 10% FCS (Sigma-Aldrich) and 100 U/ml penicillin plus 100 mg/ml streptomycin (Sigma-Aldrich) was added to the cells and cell counting was conducted by trypan blue staining. MLN cells were adjusted to 5.0 x 10^6^ cells/ml for further flow cytometric analysis.

### 2.8 Flow cytometry

MLN-derived T cell populations were phenotypically analysed as described in Myhill *et al.* (2020).^[17]^ Briefly, MLN cell suspensions were added to a 96 well plate for surface staining with anti-mouse TCRβ-FITC (clone H57-597, BD Biosciences) and anti-CD4-PerCP-Cy5.5 (clone RM4-5, BD Biosciences). Anti-T-bet-Alexafluor 647 (clone 4B10, BD Biosciences), anti-GATA3-PE (clone TWAJ, Thermo Fisher Scientific) and anti-Foxp3-FITC (clone FJK-16s, Thermo Fisher Scientific) were used for intracellular staining. MLN cell analysis was carried out on a BD Accuri C6 flow cytometer, and data acquired using Accuri CFlow Plus software (Accuri Cytometers).

### 2.9 RNA-sequencing

RNA extracted from cultured cells or caecum tissue was used for RNA-sequencing (150bp paired-end Illumina NovaSeq6000 sequencing, Novogene, Cambridge, UK). Sequence data was subsequently mapped to the *Mus muscularis* (GRCm38) genome and read counts generated which were used to determine DEG using DEseq2.^[18]^ Pathway analysis was done using the clusterProfile R package, utilizing both gene ontology and KEGG enrichment analysis. Volcano Plots were constructed using VolcaNoseR.^[19]^ Pathways with correct *P* value of <0.05 were considered significant. Sequence data is available at GEO under the accession numbers GSE178282, GSE178499, and GSE178655.

### 2.10 Quantitative real time PCR

cDNA synthesis was conducted by QuantiTect Reverse Transcription Kit (Qiagen). qPCR was performed with PerfeCTa SYBR Green FastMIX Low ROX (Quanta Bioscience). The qPCR primers are shown in Table 1. *B2m* was used as a reference gene for normalization, and fold changes calculated using the ^ΔΔ^CT method.

**Table 1.**
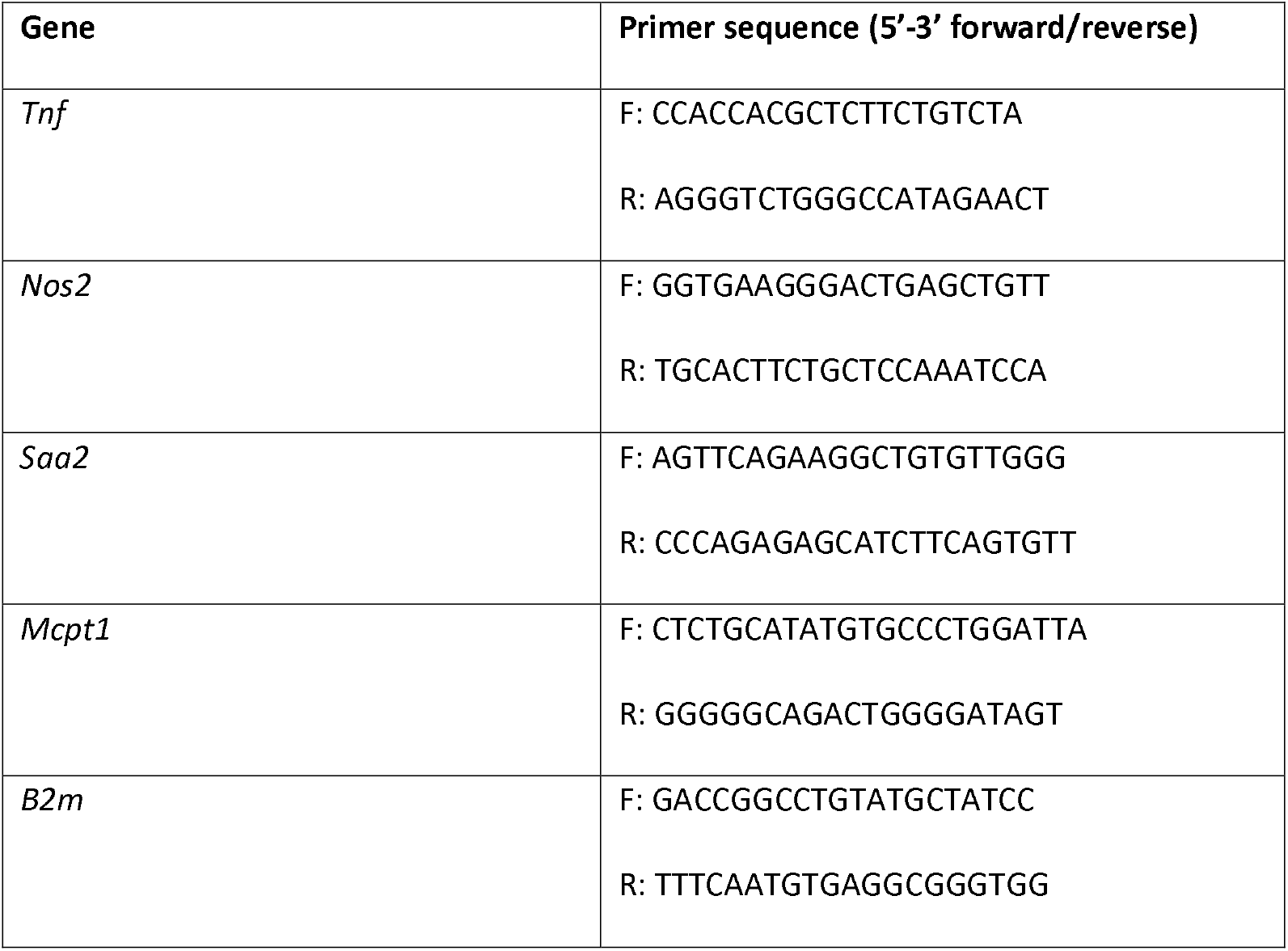
Primers used for qPCR.

### 2.11 Statistical analysis

Statistical analysis was determined by one way ANOVA with Graph Pad Prism (Version 8.0). Principal component anlysis was conducted using ClustVis.^[20]^ All data are represented as means ± standard error of mean (SEM). P value < 0.05 was considered significant.

## 3. Results

### 3.1 Effects of garlic organosulfur compounds on cytokine production and gene expression from murine macrophages

To investigate the possible anti-inflammatory effects of PTSO-PTS, we first exposed RAW264.7 macrophages to either LPS alone or LPS combined with 20 μM PTSO-PTS – the given dose was based on preliminary experiments that showed this amount was tolerated without any toxicity to cells **(Supplementary Figure** 1). After 24h culture, quantification of IL-6 and TNFα levels showed that they were markedly increased by LPS treatment alone, while concurrent PTSO-PTS exposure significantly decreased the production of these two pro-inflammatory cytokines **(Figure 1A and 1B).** Thus, PTSO-PTS has a suppressive effect on inflammatory cytokine production following toll-like receptor (TLR) stimulation.

**Figure 1.**
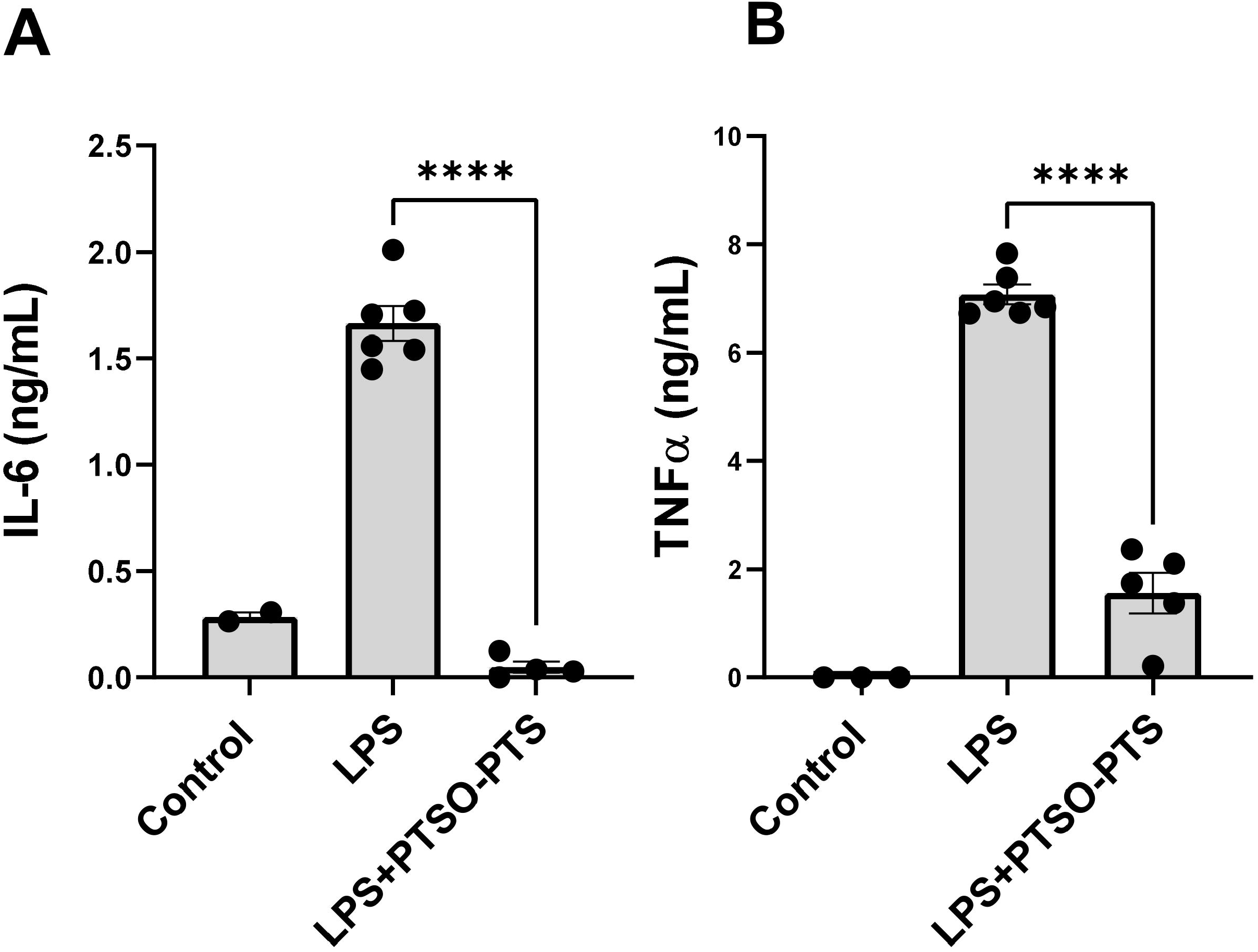
PTSO-PTS suppressed proinflammatory cytokine secretion in RAW264.7 cells. **(A)** IL-6 production in RAW264.7 cells stimulated with lipopolysaccharide (LPS) or LPS combined with 20 μM PTSO-PTS. **(B)** Cytokine TNFα in RAW264.7 cells stimulated with LPS and LPS+PTSO-PTS. Data bars represent mean + SEM. *p≤0.05, **p≤0.01, ***p≤0.001, ****p≤0.0001. Data is from at least two independent experiments.

To explore in detail the cellular activities modulated by PTSO-PTS, RNA sequencing was conducted on RAW264.7 cells following PTSO-PTS and/or LPS treatment. As shown in **Figure 2A,** in resting cells (no LPS exposure), PTSO-PTS modulated the transcription of around 700 genes (Fold change >2; adjusted *P* value <0.05). Notable upregulated genes included the transcription factors *Jun* and *Fosl2,* and the activation marker *Cd14*, suggesting that PTSO-PTS had immunogenic properties and induced activation of resting macrophages **(Figure 2B).** Downregulated genes included many involved in cell adhesion and migration, such as *Itga4, Tns3* and *Nrp2* **(Figure 2B).** Gene ontology (GO) enrichment analysis revealed that in resting cells exposed to PTSO-PTS, the main biological processes that were enriched were ubiquitin signaling, indicating an enhanced metabolic state of the cells. This was supported by KEGG pathway analysis showing that NF-kB and MAPK-related pathways, among others, were up-regulated **(Figure 2C-D).** In contrast, GO and KEGG pathway analyses demonstrated a suppression of pathways relating to cell cycle and DNA replication, as well as fatty acid metabolism **(Figure 2E-F).** Intersestingly, pathways related to cell cycle and fatty acid metabolism were also suppressed in RAW264.7 cells after exposure to LPS **(Supplementary Figure 2),** suggesting that PTSO-PTS exposure had an immunostimulatory effect on the cells. However, we could not detect any secretion of TNFα or IL-6 from PTSO-PTS treated cells without concurrent LPS exposure (data not shown).

**Figure 2.**
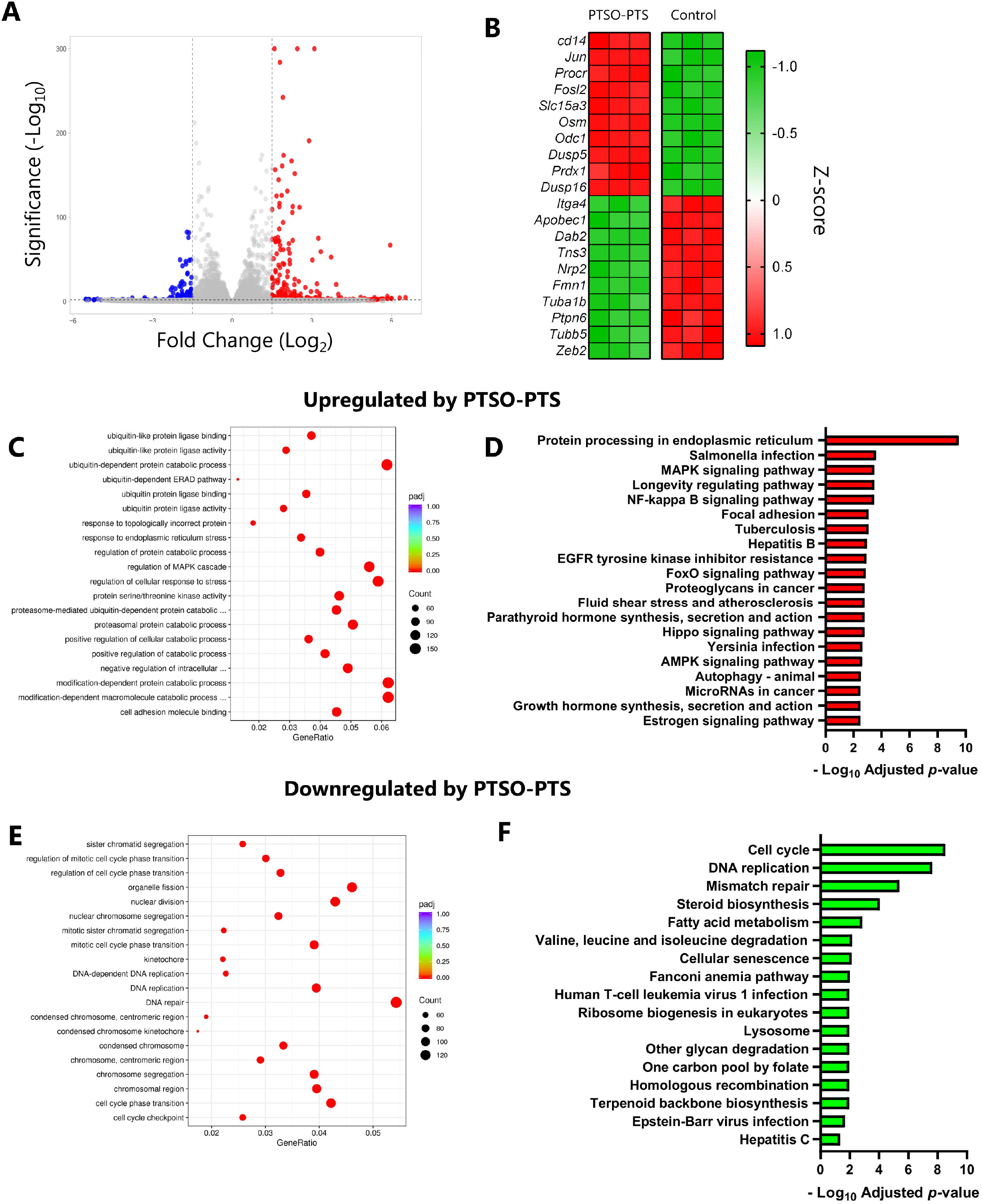
PTSO-PTS modulates transcriptional pathways in cultured RAW264.7 cells. **(A)** Volcano plot of gene expression fold changes of PTSO-PTS treated cells (20 μM for 6h) compared to non-treated controls (Fold change >2; adjusted *P* value <0.05). **(B)** Heat map of top 10 up- and down-regulated genes form PTSO-PTS treated RAW264.7 cells compared to non-treated cells, **(C)** Gene Ontology (GO) analysis of top upregulated pathways by PTSO-PTS compared to non-treatd cells. **(D)** Top upregulated KEGG pathways following exposure of cells to PTSO-PTS (padj<0.05). **(E)** GO analysis of top downregulated pathways by PTSO-PTS. **(F)** Significantly downregulated KEGG pathways following exposure of cells to PTSO-PTS (padj<0.05) n=3 per treatment group.

Given that LPS-induced cytokine production was significantly impaired by PTSO-PTS, we next investigated the transcriptomic response of LPS-activated cells with or without concurrent PTSO-PTS treatment. Around 1200 genes were significantly modulated by PTSO-PTS treatment, relative to cells exposed only to LPS **(Figure 3A).** Principal component analysis indicated that the response to LPS was profoundly altered by PTSO-PTS **(Figure 3B).** Down-regulated genes include many related to immune function, such as *Ccl22,* but also genes related to lipid metabolism *(Lpl)* and G protein-coupled receptor signaling *(Rgs2).* Notably, genes encoding inflammatory cytokines and chemokines *(Tnf, Il27, Cxcl9)* were suppressed. However, we also noted that other cytokine-encoding genes (e.g. *Il23a, Csf3)* were increased in LPS-treated cells exposed to PTSO-PTS, suggesting a pleiotropic effect **(Figure 3A and C).** Interestingly, as observed in resting cells, expression of *Cd14, Procr* and *Osm* was significantly increased by PTSO-PTS in LPS-activated cells, and GO analysis again revealed that a dominant biological response to PTSO-PTS was ubiquitin activity **(Figure 3D).** KEGG pathway analysis showed many enriched pathways related to glutathione and lysosome activity, suggesting that xenobiotic metabolizing pathways were activated in response to PTSO-PTS exposure **(Figure 3E).** Downregulated pathways identified by GO and KEGG analysis mainly related to immune defense and interferon signaling, consistent with the previously observed anti-inflammatory activity **(Figure 3F-G).** To verify the anti-inflammatory pathways identified in our transcriptomic analysis, we confirmed that LPS-induced IL-27 secretion was significantly impaired by PTSO-PTS **(Figure 3H).** Collectively, these data show that resting macrophages adopt a semi-activated state after exposure to PTSO-PTS, but that TLR4-induced inflammatory pathways are strongly suppressed, indicating that PTSO-PTS is likely to demonstrate profound immunomodulatory properties in inflammatory settings.

**Figure 3.**
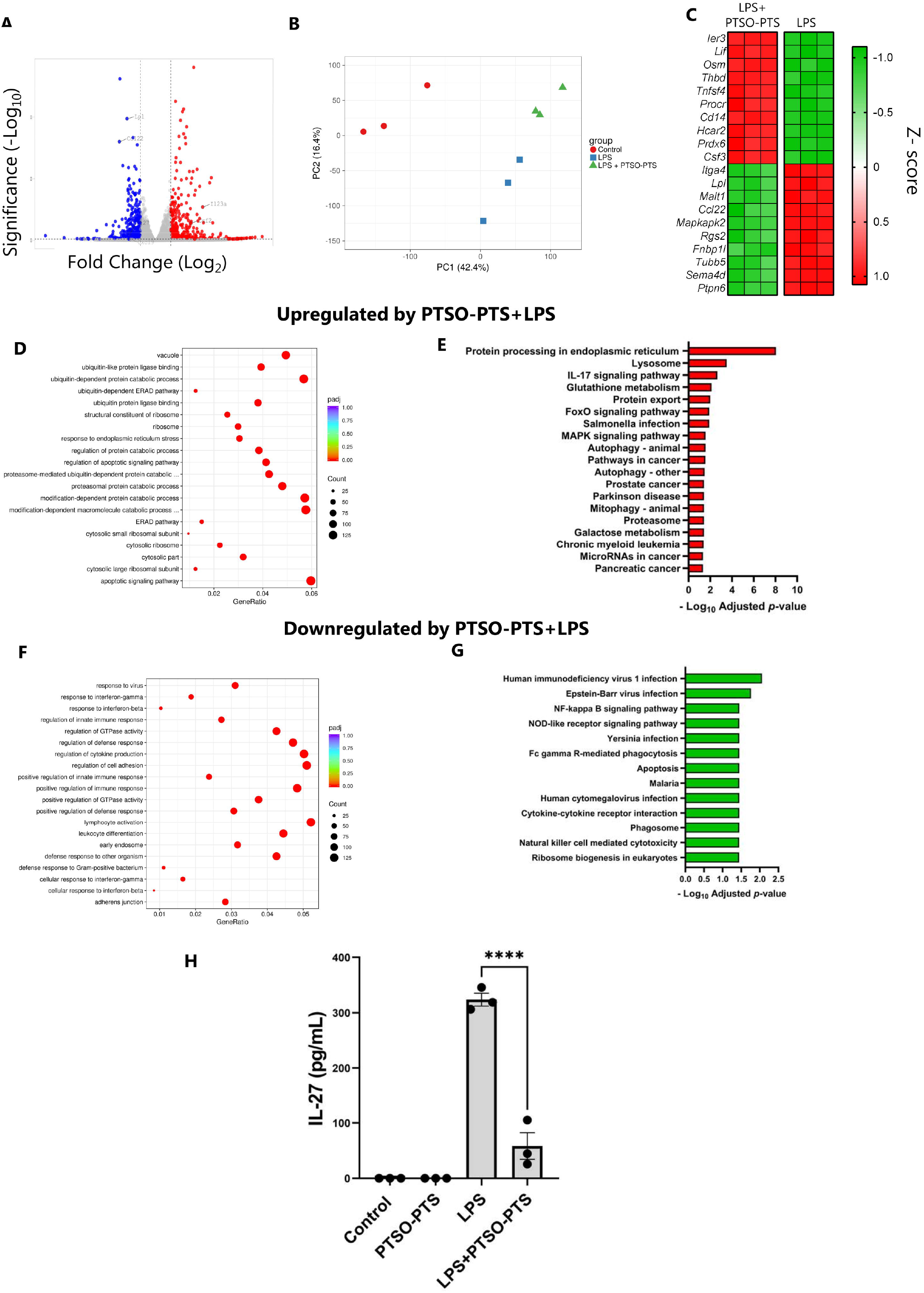
PTSO-PTS regulates transcriptional pathways and inflammatory responses following lipopolysaccharide treatment in RAW264.7 cells. **(A)** Volcano plot of genes expressed showing fold changes in PTSO-PTS and lipopolysaccharide (LPS) treated cells compared to LPS-only treated controls. **(B)** Principal component analysis showing clustering of cells without stimulation, LPS stimulation, or LPS combined with 20 μM PTSO-PTS **(C)** Heat map of top 10 up- and down-regulated genes in PTSO-PTS +LPS treated RAW264.7 cells compared to only LPS treated cells. **(D)** Gene ontology (GO) analysis of top upregulated pathways in LPS-treated cells exposed to PTSO-PTS compared to LPS-only treated controls. **(E)** Top upregulated KEGG pathways in LPS-treated cells following exposure to PTSO-PTS, compared to LPS-only treated controls (padj<0.05). **(F)** GO analysis of top downregulated pathways in LPS-treated cells exposed to PTSO-PTS compared to LPS-only treated controls. **(G)** Significantly upregulated KEGG pathways in LPS-treated cells following exposure to PTSO-PTS, compared to LPS-only treated controls (padj<0.05). **(H)** IL-27 secretion from LPS-stimulated RAW264.7 macrophages cultured with PTSO-PTS. Data bars represent mean + SEM. ****p≤0.0001. n=3 per treatment group.

### 3.2 Effects of PTSO-PTS treatment on murine intestinal epithelial cells

As dietary components, the garlic-derived compounds likely come into contact with the intestinal epithelium in relatively high concentrations before and during absorption. Therefore, we also investigated whether PTSO-PTS modulated the activity of cultured intestinal epithelial cells (IEC) by exposing CMT-93 cells to PTSO-PTS for 6 hours, followed by RNA sequencing. The dominant transcriptional response to PTSO-PTS was related to cell cycle and DNA replication activity **(Figure 4A-B).** Moreover, consistent with the indications of xenobiotic metabolism-related transcriptional responses in macrophages, the most significantly up-regulated genes were involved in antioxidant responses *(Gpx2,* and the selenoprotein-encoding genes *Selenow* and *Selenoh;* **Figure 4C).** Thus, PTSO-PTS has a stimulatory effect on IEC under steady-state conditions, and induces genes related to detoxification and antioxidant activity. To induce an inflammatory response in these cells, we exposed them to antigens from an enteric pathogen. *T. muris* causes chronic infections with colitis-like inflammation, and its secreted antigens have been shown to induce the expression of pro-inflammatory genes in IEC *in vitro.*^[21]^ Thus, to explore the effect of PTSO-PTS on IEC responses during exposure to pathogen antigens, we conducted RNA sequencing on *T. muris* excretory/secretory antigen (E/S)-stimulated cells with or without concurrent PTSO-PTS treatment. CMT-93 cells reacted to *T. muris* E/S by up-regulating transcriptional pathways related to cellular activity such as DNA replication and ribosome activity, and genes encoding nutrient transporters *(Slc7a11, Tfrc)* and cytokines/chemokines (*Il24, Cxc15*) **(Figure 4D-F).** PTSO-PTS-induced transcriptional changes in cell cycle pathways were maintained during exposure to *T. muris* E/S. These included MAPK signaling and cell cycle pathways **(Figure 4H).** Expression of the antioxidant-related genes *Gpx1* and *Selenow* was also upregulated by PTSO-PTS in *T. muris* E/S-treated cells **(Figure 4I).** These data show that PTSO-PTS treatment induced a varied transcriptomic response in IEC both alone and also in the presence of pathogen antigens, suggesting that PTSO-PTS may exert immunomodulatory effects by contacting intestinal epithelium during enteric infection and inflammation.

**Figure 4.**
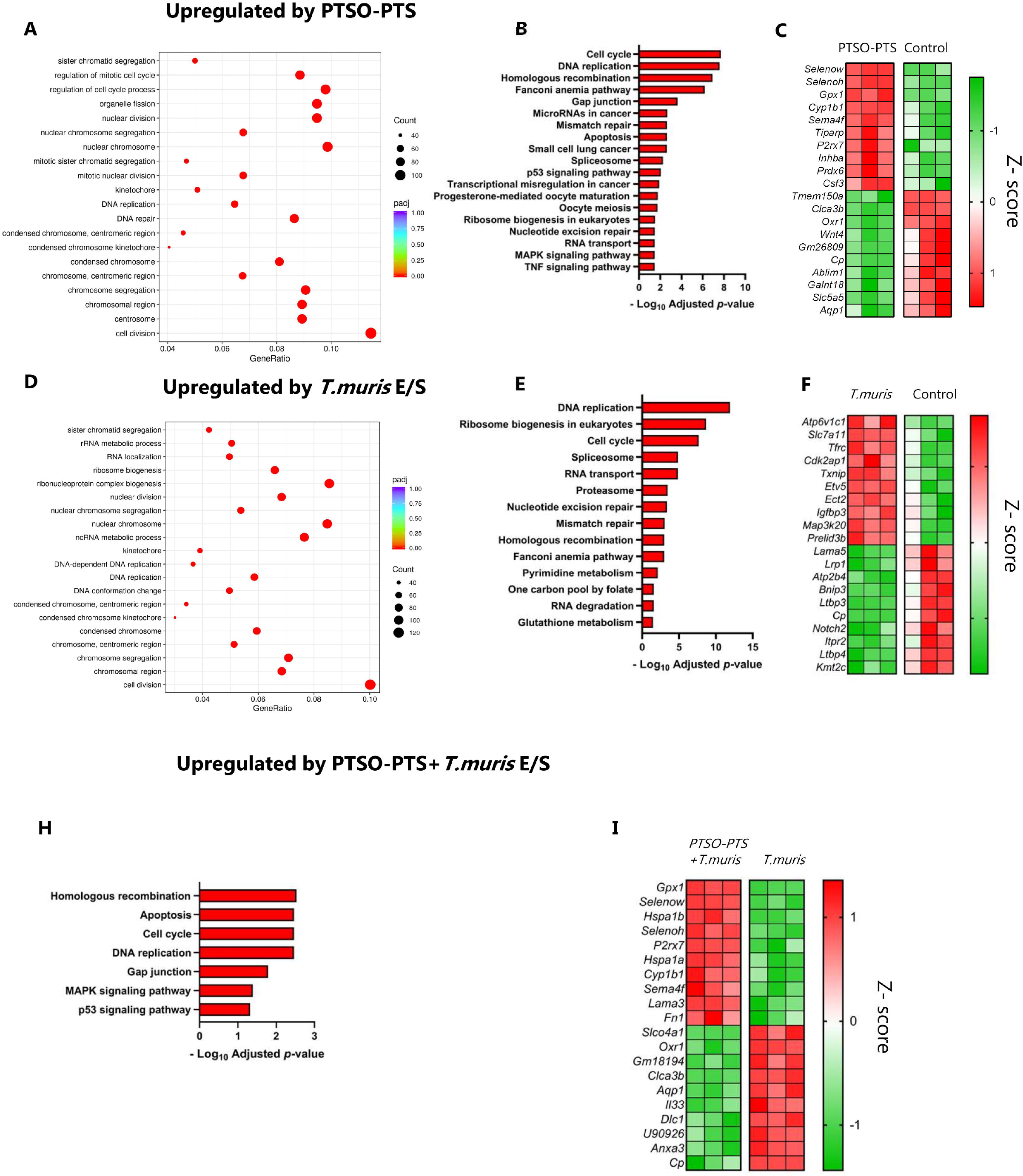
PTSO-PTS modulates transcriptional pathways in cultured intestinal epithelial cells during *Tríchurís muris* antigen stimulation. **(A)** Gene ontology (GO) analysis of top upregulated pathways by PTSO-PTS in CMT93 cells, compared to untreated cells. **(B)** KEGG pathways upregulated following exposure of CMT93 cells to PTSO-PTS, compared to untreated cells (padj<0.05). **(C)** Heat map of top 10 up- and down-regulated genes in CMT93 cells treated PTSO-PTS, compared to untreated cells, **(D)** GO analysis of top upregulated pathways following exposure of cells to *T.muris* E/S antigens, compared to untreated cells. **(E)** KEGG pathways significantly upregulated by *T.muris* antigens in CMT93 cells, compared to untreated cells (padj<0.05). **(F)** Heat map of top up- and down-regulated genes by *T.muris* antigens, compared to untreated cells **(G)** KEGG pathways upregulated in *T.muris* antigen-treated cells exposed to by PTSO-PTS, relative to only *T.muris* antigen treated cells (padj<0.05). **(H)** Heat map of top up- and down-regulated genes in *T.muris* antigen-treated cells exposed to by PTSO-PTS, relative to only *T.muris* antigen treated cells n=3 per treatment group.

### 3.3 PTSO-PTS does not affect *Trichuris muris* burdens but modulates expression of inflammation-related genes

We next asked if oral treatment of mice with PTSO-PTS could regulate immune function and inflammatory responses during *T. muris* infection. In C57BL/6 mice, infection with low numbers of *T. muris* eggs (≤ 20) causes a chronic infection in the caecum, which is associated with the expression of numerous inflammation-related genes and the development of sporadic colitis like-pathology.^[22,23]^ The ability of PTSO-PTS to potentially reduce enteric infection and inflammation *in vivo* was studied by daily feeding 0.1 mg/kg BW PTSO-PTS and 1 mg/kg BW PTSO-PTS to mice during a chronic *T. muris* infection. For both concentrations of PTSO-PTS, there was no effect on worm burdens compared to *T. muris*-infected mice alone **(Figure 5A).** Although mice fed PTSO-PTS had no changes in daily weight gain during the entire experiment, *T. muris*-infected mice administered 1 mg/kg PTSO-PTS displayed decreased weight gain compared to untreated, infected mice during infection **(Figure 5B and 5C,** respectively).

**Figure 5.**
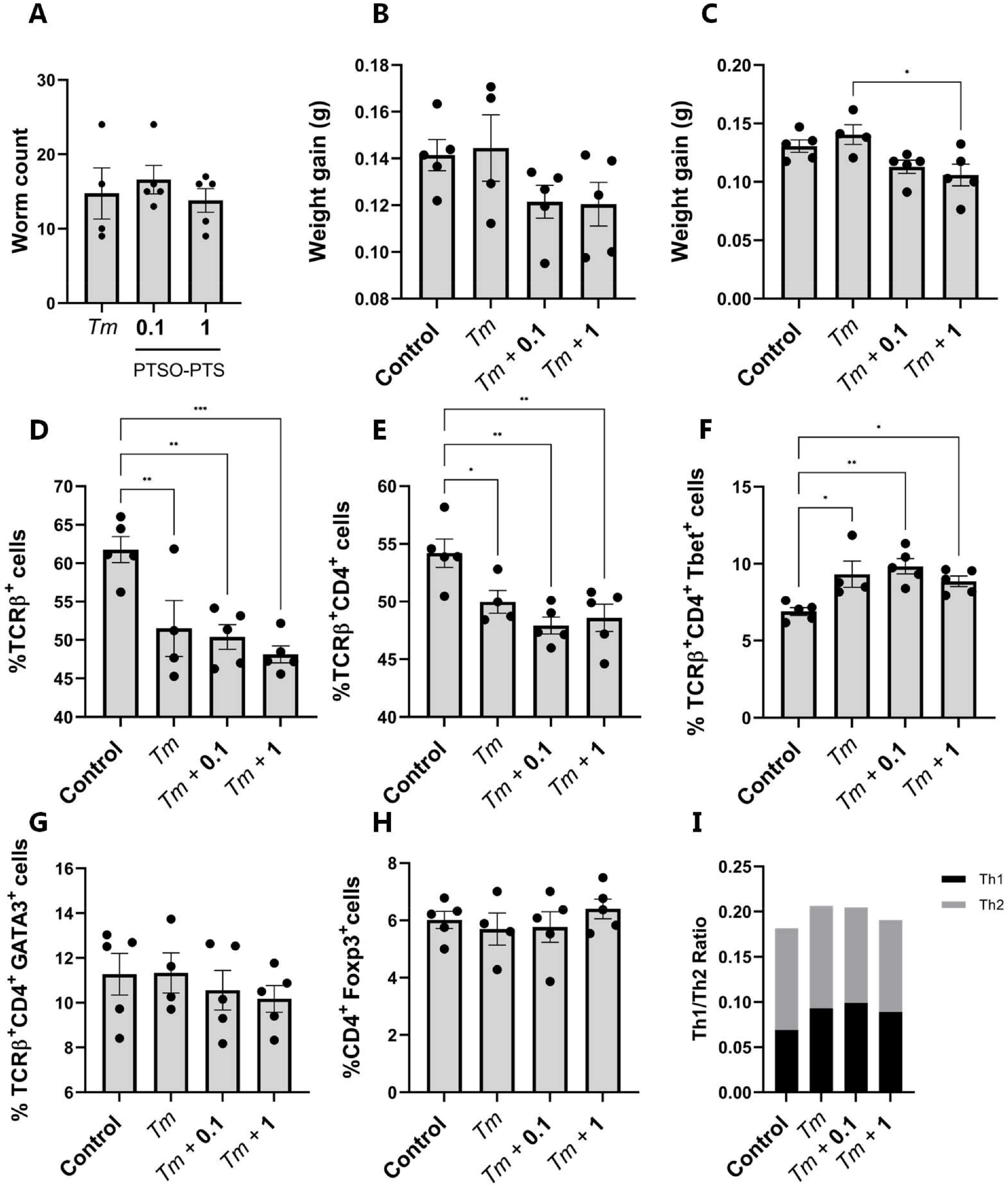
*Tríchurís muris* infection regulates immune responses from mesenteric lymph node cells rather than PTSO-PTS. **(A)** *T. muris* burdens in mice inoculated with 20 eggs and given control drinking water, or water supplemented with either 0.1 mg/kg or 1 mg/kg PTSO-PTS. Worm burdens were assessed at day 35 post-infection. Weight gain throughout the entire experimental period **(B)** and during infection with *T. muris* **(C). D)** Proportions of TCRβ^+^ cells in mesenteric lymph nodes (MLN) following 35 days of *T. muris* infection in mice given control drinking water, or water supplemented with either 0.1 mg/kg or 1 mg/kg PTSO-PTS. Shown also are uninfected control mice given normal drinking water. Proportions of CD4 ^+^ (E), CD4^+^T-bet^+^**(F),** CD4^+^GATA3^+^**(G)** and CD4^+^Foxp3^+^T-cells in MLN following 35 days of *T. muris* infection in mice given control drinking water, or water supplemented with either 0.1 mg/kg or 1 mg/kg PTSO-PTS. Shown also are uninfected control mice given normal drinking water. I) Th2 (CD4^+^GATA3^+^) to Th1 (CD4^+^T-bef) ratio in MLN. * p<0.05, ** p<0.01, **p<0.001 by ANOVA analysis, n= 4-5 per treatment group.

To explore the effect of PTSO-PTS on host immune responses, mesenteric lymph node (MLN) cells were isolated for flow cytometric analysis. As shown in Figure **5D** and **5E**, *T. muris* infection clearly reduced the proportion of TCRβ^+^ T cells and TCRβ^+^CD4^+^ T cells, but there was no effect of PTSO-PTS intake on the proportions of these cells within infected mice. Within the T-cell population there was a significant increase in the proportion of Th1 cells (TCRβ^+^CD4^+^Tbet^+^) in infected mice, consistent with the type-1 immune response known to be induced by chronic *T. muris* infection, but again PTSO-PTS intake did not influence these parameters **(Figure 5F).** The proportions of Th2 (GATA3^+^) and T-regulatory (Foxp3^+^) T-cells were not altered by either infection or PTSO-PTS intake **(Figure 5G, 5H and 5I).**

To explore in more detail whether PTSO-PTS could modulate the mucosal inflammation induced by the infection, RNA sequencing was carried out on caecal tissue harvested from mice in each treatment group. As expected, *T. muris* infection altered the expression of a vast number of genes (>3500), relative to uninfected control mice. Amongst the top up-regulated genes were the granzyme-encoding genes *Gzma* and *Gzmb,* the interferon-responsive gene *Ifit2,* and the inflammatory chemokine *Ccl5* **(Figure 6A).** Consistent with this, enriched KEGG pathways were related to infection and inflammation, including TNF signaling **(Supplementary Figure 3).** Interestingly, numerous KEGG pathways related to nutrient metabolism (e.g. amino acids, fatty acid metabolism) were suppressed by infection **(Supplementary Figure 4).** Principal component analysis demonstrated that, within infected mice, PTSO-PTS intake resulted in dose-dependence divergence from control mice **(Figure 6B).** However, only a small number of genes were identified as being significantly (adjusted *p* value <0.05) regulated by PTSO-PTS within infected mice. These included *Upk3b* (encoding an epithelial protein found in the intestinal and urogenital tracts), *Lrrn4* (encoding a leucine-rich protein involved in diverse cellular functions), and *Myrf* (encoding a myelin regulatory factor) – these genes were upregulated in both treatment groups (0.1 mg/kg and 1 mg/kg). In the higher dosage group, an additional three genes were regulated. These genes encoded a glutamate transporter *(Slc17a8),* an insulin-like growth factor binding protein *(Igfbp6),* and an uncharacterized immunoglobulin variable region gene *(Igkv5-37). We* also noted that both PTSO-PTS treated groups induced a significant (adjusted *p* value <0.05) down-regulation of several KEGG pathways related to protein metabolism, inflammation and immune function. Interestingly, these pathways tended to be more strongly regulated in the 0.1 mg/kg group **(Figure 6C-D).** Further examination of the transcriptomic data showed that a number of genes related to inflammation (including *Tnf, Nos2* and *Saa2)* were down-regulated by PTSO-PTS. Conversely, genes related to the development of type-2 immunity (e.g. *Mcpt1*), were upregulated. As these genes were not significant after correcting for multiple testing, we verified their expression by qPCR. This confirmed a significant dose-dependent suppression of *Tnf, (Nos2)* and *Saa2* expression (and an increase in *Mcpt1* expression) **(Figure 6E).** Thus, in this model PTSO-PTS intake was not sufficient to lower parasite burdens but did appear to attenuate the localized pro-inflammatory transcriptional response in the caecum.

**Figure 6.**
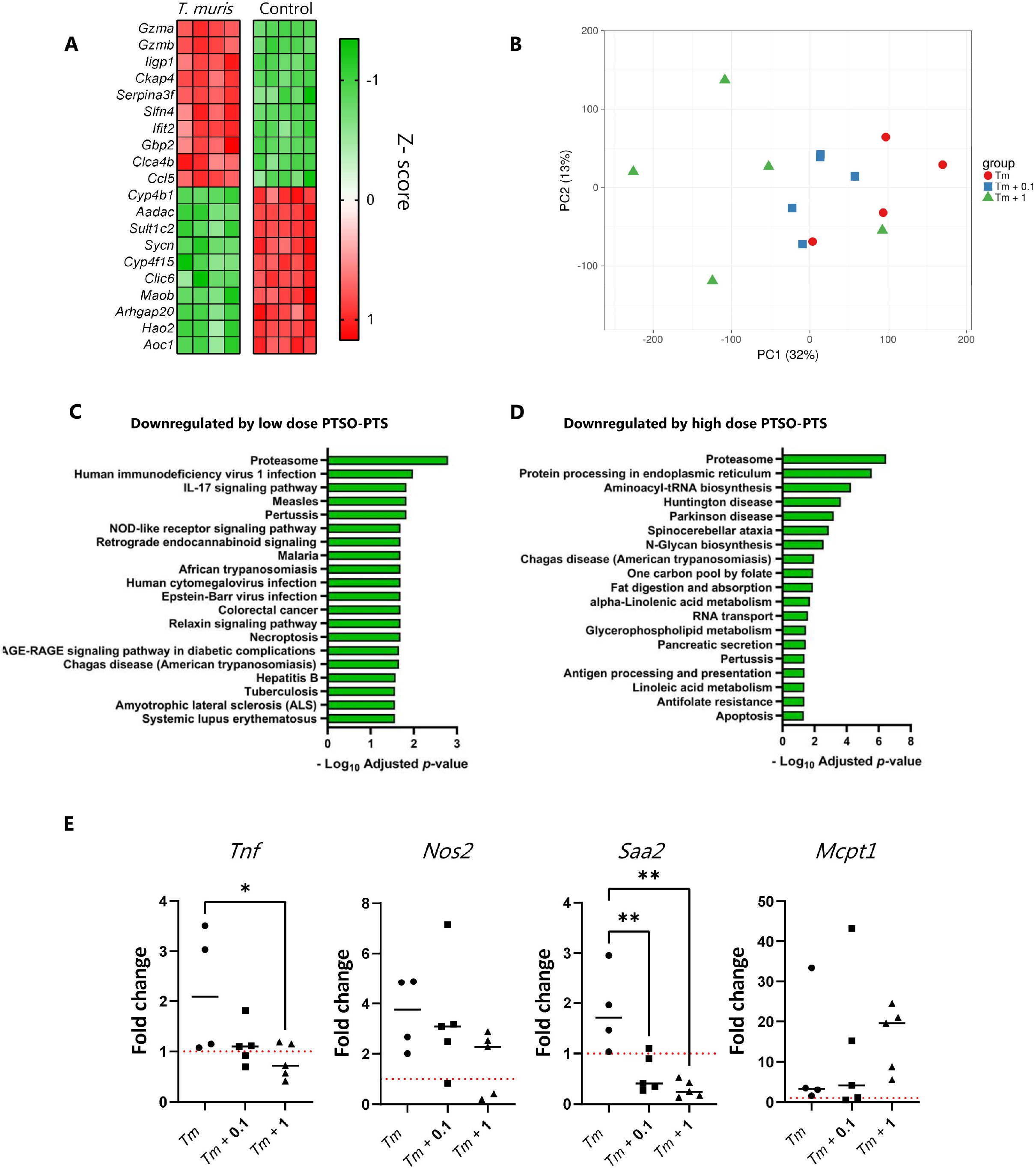
PTSO-PTS modulates the pro-inflammatory transcriptional response in the mouse caecum. **(A)** Top ten most up- and down-regulated genes in caecal tissue of *Trichuris muris*-infected mice, relative to control, uninfected mice. **B)** Principal component analysis showing clustering of caecal tissue in uninfected mice, and mice infected with *T. muris* for 35 days and given control drinking water, or water supplemented with either 0.1 mg/kg or 1 mg/kg PTSO-PTS. KEGG biological pathways significantly down-regulated in caecal tissue of *T. muris*-infected mice given either 0.1 **(C)** mg/kg or 1 mg/kg **(D)** PTSO-PTS, relative to *T. muris*-infected mice given control drinking water. **E)** qPCR validation of the expression of *Tnf Nos2, Saa2* and *Mcpt1* in caecal tissue following 35 days of *T. muris* infection in mice given control drinking water, or water supplemented with either 0.1 mg/kg or 1 mg/kg PTSO-PTS. Shown also are uninfected control mice given normal drinking water. * p<0.05, ** p<0.01, by ANOVA analysis, n= 4-5 per treatment group.

## 4 Discussion

In recent years, garlic-derived compounds have been increasingly studied as a therapeutic agent. They have characteristic chemical structures, especially sulfur containing compounds, sulfides and alliin, which may influence their biological activities to protect animals and humans from diseases.^[24]^ In most compounds of garlic, PTSO-PTS show pungent odors, and PTS is produced by the interaction of alliinase and S-propyl-l-cysteine sulfoxide, and then changes into PTSO due to its instability.^[25]^ PTSO-PTS is absorbed mainly in the intestine by amino acid transporter of cysteine, showing broad antibiotic activity.^[24,26]^ Interestingly, PTSO-PTS has also demonstrated immunomodulatory properties and can alleviate localized inflammation.^[13]^ During transit throughout the digestive tract, PTSO-PTS may modify intestinal responses by contacting intestinal cells, and we thus explored the PTSO-PTS effects on murine intestine *in vitro* and *in vivo* to elucidate the underlying anti-inflammatory mechanisms and whether these compounds may have protective effects during chroninc enteric infection.

Murine RAW264.7 macrophages are well-known to secrete pro-inflammatory mediators and thus mimick inflammation status during LPS activation.^[27]^ We initially observed that PTSO-PTS induced dramatic changes by inhibiting the secretion of the pro-inflammatory cytokines TNF-α and IL-6 in RAW 264.7 cells, indicating potential anti-inflammatory ability. In addition, the gene expression profile of macrophages was markedsly affected by PTSO-PTS treatment. Interestingly, PTSO-PTS induced functional markers *Cd14, Procr* and *Osm* in resting cells, showing that PTSO-PTS may have an immune-activation effect.^[28],[29]^ Notably, the suppressed pathways mainly refer to DNA replication and cell cycle, which is consistent with anticancer properties of garlic-derived extracts.^[30]^

During exposure to LPS, PTSO-PTS suppressed the induction of inflammatory markers in RAW264.7 cells, such as *Ccl22,* a chemokine related to cell migration and inflammatory function which can be induced to promote tissue damage, and is a main factor of pathogenesis of asthma and allergy.^[31]^ Moreover, anti-*Ccl22* treatment has the ability to ameliorate CNS inflammation by modulating inflammatory macrophage accumulation and macrophage effector cytokine phenotype.^[32]^ Genes encoding *Il27*, a cytokine belonging to the IL-6 family, and *Cxcl9,* a chemokine involved in inflammation and tumor progression, were both significantly downregulated in LPS-induced macrophages, confirming the strong anti-inflammatory effect of PTSO-PTS.^[33,34]^ Interestingly, we noted that the glutathione activity was upregulated by PTSO-PTS, which is related to oxidative response and detoxifying effects as shown by Anoush *et al* ^[35]^ using garlic extracts, suggesting PTSO-PTS is involved in detoxification and xenobiotic elimination. Taken together, PTSO-PTS seems to exert immunostimulatory effects in resting cells and anti-inflammatory properties towards TLR-4 activated responses induced by LPS.

Allicin has been reported to produce metabolites that can contact and interact with intestinal cells.^[24]^ However, whether PTSO-PTS plays a similar role via contacting intestinal epithelial cells is unclear. The RNA-seq of CMT cells with PTSO-PTS indicates regulation of antioxidant responses. *Gpx1* is associated with endothelial dysfunction.^[36]^ Upregulation of *Gpx1* is known to provide protection of ileum and colon mucosa from inflammation and oxidative stress with enhanced genes *Selenow* and *Selenoh,* which belong to intracellular selenium (Se)-dependent glutathione peroxidase, contributing to glutathione activity as similarly observed in RAW macrophages.^[37]^ Notably, apoptosis pathway induced by PTSO-PTS is consistent with the garlic compound ajoene which has been shown to limit tumor size in cancer.^[38]^

*T. muris* E/S products contain multiple protein and small molecules secreted by *T. muris,* which are known to regulate host immunity.^[39]^ Similarly, we found that treatment with *T. muris* E/S alone *in vitro* also tends to regulate energy metabolism including glycolysis, galactose metabolism and glucagon signaling as previously reported.^[40]^ PTSO-PTS treatment with *T. muris* E/S modulated a variety of intestinal transcriptional responses. The significantly up-regulated pathways were DNA replication and anabolic pathway, which were down-regulated in macrophages, indicating PTSO-PTS exerts pleiotropic effects in various cell lines. Furthermore, the significantly regulated genes *Gpx1* and *Selenow* were also observed. Thus, our data suggests that antioxidant responses in intestinal epithelial cells may be important mechanism by which PTSO-PTS to limit infection.

Low dose *T. muris* inoculation in mice tends to drive a chronic infection, characterized by Th1 immune polarization, and intestinal inflammation.^[41]^ Garlic has exhibited anticoccidial activity by impairing the development of parasites in a murine model, demonstrating the possibility of garlic as an anti-parasite agent (in a Th1 dominated inflammatory state).^[42]^ Our *in vivo* study showed that PTSO-PTS supplement did not alter *T. muris* worm counts, which is in contrast to garlic extracts,^[43]^ showing that PTSO-PTS may not be a suitable anti-parasite compond against *T.muris,* yet additional experiments are needed to fully investigate the effects of PTSO-PTS on intestinal parasitic worms. Furthermore, we observed a distinct Th1-polarized immune response towards *T. muris,* yet no changes in Th1, Th2 or Treg cell populations were induced by PTSO-PTS treatment during *T. muris* infection. Thus, our data confirms that chronic *T. muris* infection causes the polarization of a type-1 immune response as previously reported,^[44]^ and PTSO-PTS has no effect on MLN-derived CD4^+^T cell phenotypes.

In mice infected with low levels of *T. muris,* infection resulted in host defence transcriptional responses, especially upregulated T cell differentiation, TLR and TNF signaling, and interferon gamma response as reported in other studies.^[45]^ Interestingly, *T. muris* infection suppressed metabolism of xenobiotics by cytochrome p450 response. The downregulation of cytochrome p450 reflects the host resistance mechanism against intestinal parasites, further supporting oxidative stress activity and pro-inflammation of parasite.^[46]^ Similarly, PTSO-PTS treatment in infected mice induced significant changes to intestinal gene expression and pathways related to immune function and inflammation, which is consistent with protective effect of PTSO-PTS against avian coccidiosis.^[47]^ It is interesting to note the downregulated IL17 signaling pathway in 0.1 mg/kg PTSO-PTS dosed mice rather than higher PTSO-PTS concentration, as uncontrolled IL17 signaling can exacerbate autoimmune diseases and immunopathology.^[48]^ Thus, the anti-inflammatory property of PTSO-PTS seems to be related to concentration, which could be investigated further in relation to treating autoimmune pathologies. During inflammation, the intestinal barrier is damaged as a result of excessive reactive nitrogen and oxygen (NO) species production, induced by the enzyme NOS2.^[49]^ The NOS2-induced production of NO can be stimulated by pro-inflammatory cytokines, such as TNFα and IFN-γ, concomitant with selenoprotein inhibition [49]. Serum amyloid A2, SAA2, is activated during infection and tissue injury as a marker of inflammation, and is involved in various chronic diseases.^[50]^ Mucosal mast cell proteases (e.g. *Mcpt1*) are known to be induced by TGF-β, which may be involved in degradation of pro-inflammatory substances and thus exerts anti-inflammation activity.^[51],[52]^ Taken together, the downregulation of *Nos2, Saa2* and upregulation of *Mcpt1* further confirms the anti-inflammatory ability of PTSO-PTS during *T. muris* infection. The anti-inflammatory activity of PTSO-PTS in this model may derive from the antioxidant acitivty induced in immune cells at the mucosal barrier surface, as evidenced by our *in vitro* data, although future studies should also assess whether these compounds may interact with the gut microbiota to exert immunnomodlulatory effects, e.g. by the production of metabolites with systemic antiinflammatory properties.

This study clearly demonstrates the anti-oxidant and anti-inflammatory ability of PTSO-PTS *in vitro* and *in vivo,* with these properties further demonstrated by the regulation of the gut environment resulting from the beneficial interplay between diet and helminth infection. Our results encourage further investigations to explore how bioactive compounds derived from common food sources may be used to prevent and/or treat enteric diseases.

## Abbreviations

E/S: Excretory/secretory antigen
FCS: Fetal calf serum
IEC: Intestinal epithelial cells
LPS: Lipopolysaccharide
MAPK: Mitogen-activated protein kinase
MLN: Mesenteric lymph node
Myrf: Myelin regulatory factor
PTS: Propyl-propane thiosulfinate
PTSO: Propyl-propane thiosulfonate
Th: T-helper cell type
TLR: Toll-like receptors;

## Author Contributions

LZ and ARW designed the study. LZ, LJM and AAC performed the experiments. SMT, AB and ARW supervised the study and provided essential reagents. LZ and ARW analysed the data and wrote the paper. All authors reviewed and edited the final manuscript.

## Acknowledgements

This work was supported by Independent Research Fund Denmark (Grant 7026-0094B) and the Novo Nordisk Foundation (Grant 0052422). LZ was supported by the China Scholarship Council (Grant 201806910065).

