## Supplementary Information for "Garlic-derived organosulfur compounds regulate metabolic and immune pathways in macrophages and attenuate intestinal inflammation in mice"

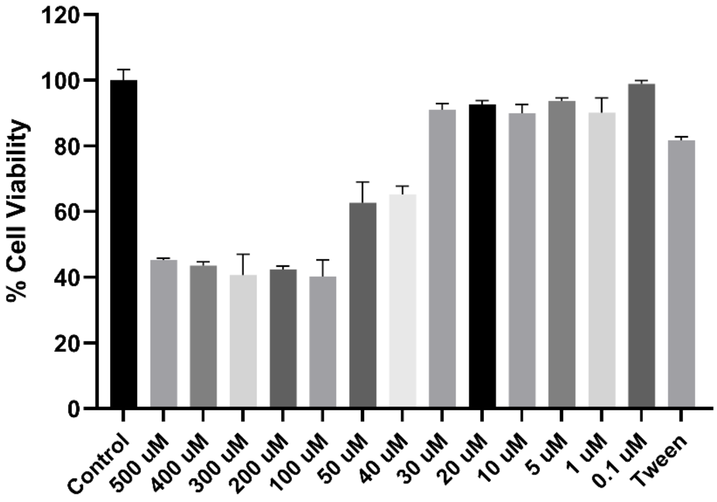


**Supplementary Figure 1:**

Cell viability of PTSO-PTS treated RAW264.7 macrophages.

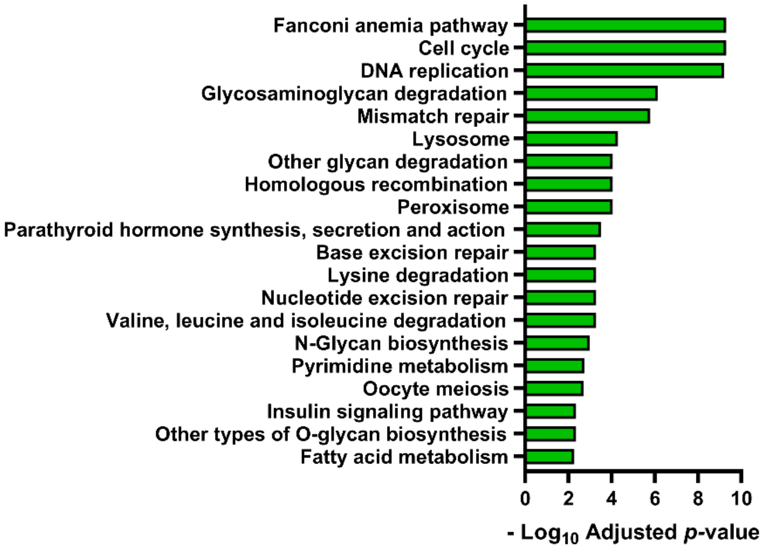


**Supplementary Figure 2.** KEGG pathways downregulated by LPS treatment in RAW264.7 cells (*p*adj<0.05), relative to untreated cells (n=3).

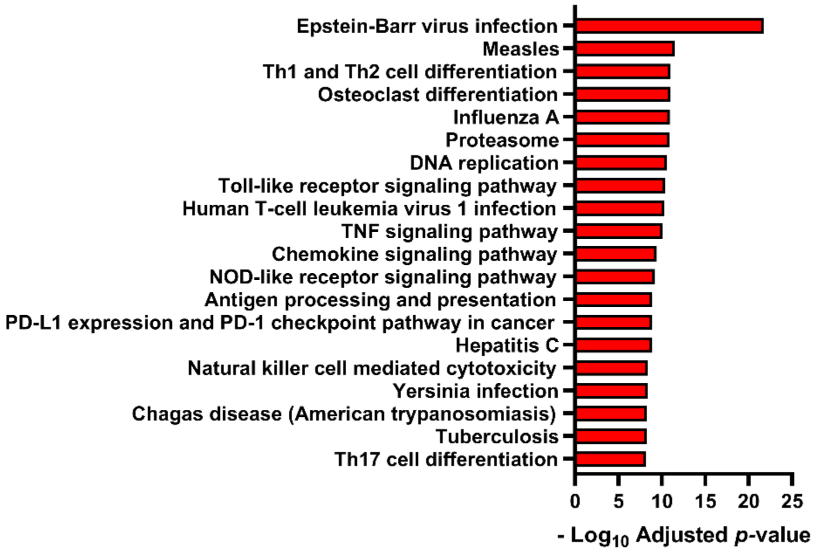


**Supplementary Figure 3.** KEGG pathways upregulated in caecal tissue by *Trichuris muris* infection , compared to uninfected mice (*p*adj<0.05).

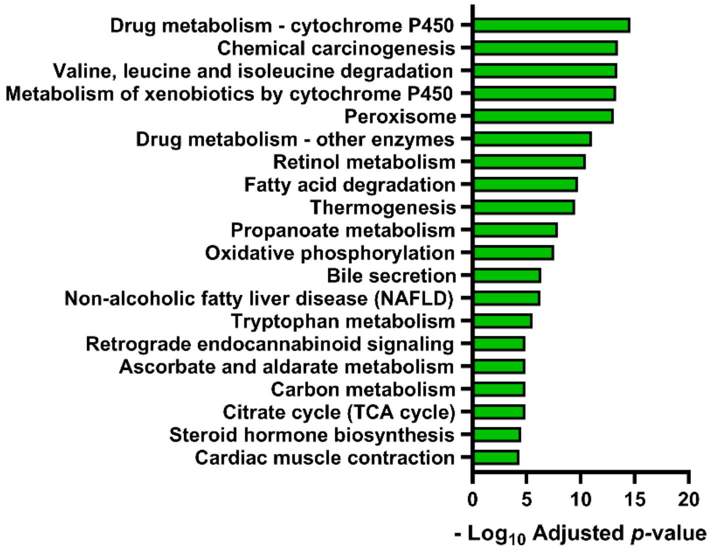


**Supplementary Figure 4.** KEGG pathways upregulated in caecal tissue by *Trichuris muris* infection , compared to uninfected mice (*p*adj<0.05).
